# The natural sequence of events in larval settlement and metamorphosis of *Hydroides elegans* (Polychaeta; Serpulidae)

**DOI:** 10.1101/2021.03.24.436767

**Authors:** Michael G. Hadfield, Marnie L. Freckelton, Brian T. Nedved

## Abstract

The broadly distributed serpulid worm *Hydroides elegans* has become a model organism for studies of marine biofouling, development and the processes of larval settlement and metamorphosis induced by surface microbial films. Contrasting descriptions of the initial events of these recruitment processes, whether settlement is induced by (1) natural multi-species biofilms, (2) biofilms composed of single bacterial species known to induce settlement, or (3) a bacterial extract stimulated the research described here. We found that settlement induced by natural biofilms or biofilms formed by the bacterium *Pseudoalteromonas luteoviolacea* is invariably initiated by attachment and secretion of an adherent and larva-enveloping primary tube, followed by loss of motile cilia and ciliated cells and morphogenesis. The bacterial extract containing complex tailocin arrays derived from an assemblage of phage genes incorporated into the bacterial genome appears to induce settlement events by destruction of larval cilia and ciliated cells, followed by attachment and primary-tube formation. Similar destruction occurred when precompetent larvae and larvae of a nudibranch gastropod were exposed to the extract, neither of which metamorphosed. We further argue that larvae that lose their cilia before attachment would be swept away from the sites that stimulated settlement by the turbulent flow characteristic of most marine habitats.

## Introduction

The life cycle of the warm-water, marine, biofouling, serpulid polychaete *Hydroides elegans* (Haswell, 1883) has been described in many publications [e.g., 1,2] (Fig. 1). Sessile adult worms live in bays and estuaries attached to rocks, mangrove roots and, problematically, the hulls of ships and the pilings where they dock. Separate sexes spawn gametes into the surrounding waters where fertilization and larval development take place. At water temperatures about 25°C, larvae begin feeding 12 h after fertilization and are metamorphically competent on day 5 [2]. Settlement and metamorphosis readily progress when, and only when, competent larvae physically contact a microbially biofilmed marine surface [3–5].

**Fig. 1.**
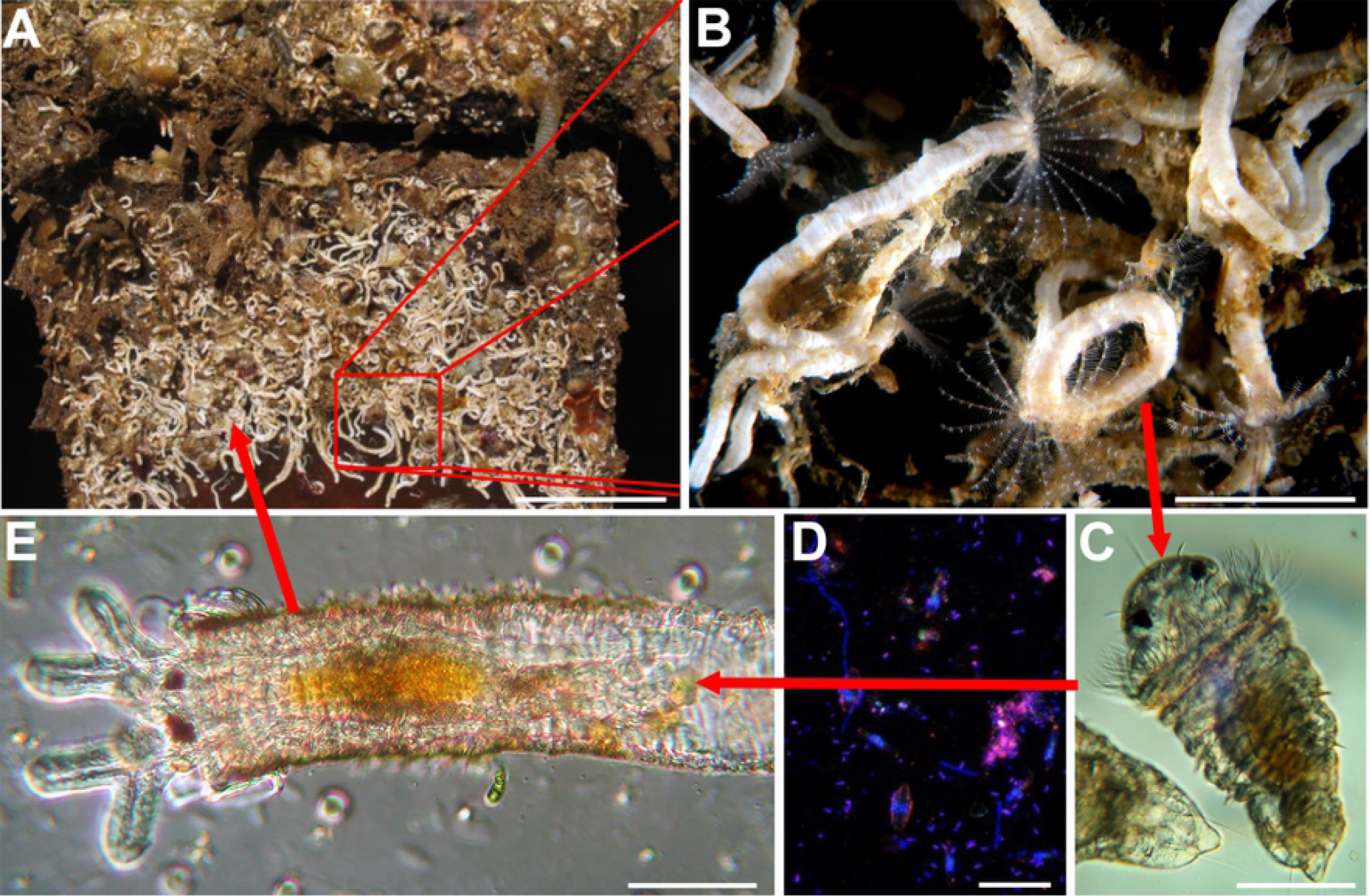
The life cycle of *H. elegans*. (A) Mature population of *H. elegans*. (B) Close up of an adult *H. elegans*. (C) competent larva of *H. elegans*. (D) biofilm bacterial surface required for larval metamorphosis. (E) early juvenile of *H. elegans*. Scale bars: A, 5cm; B, 1 cm; C, 50μm; D, 5 μm; E, 50 μm.

In the past 25 years, much of the settlement and metamorphosis processes of *H. elegans* has been substantiated in both our and other laboratories. It is known, for example, that settlement of competent larvae increases with increasing bacterial density on a surface, and that, in a natural biofilm, settlement is cued by many different, but not all, bacterial species [3,4,6,7]. Further, larvae must physically contact a biofilmed surface to perceive the bacterial cue to settle. Larvae exhibit no settlement behavior when separated from a biofilmed surface by as few as 20 μm; i.e., **there is no soluble settlement cue from bacteria in a natural biofilm** [8].

Of the marine bacteria that induce larvae of *H. elegans* to settle, one that has been intensely studied is the globally distributed, gram-negative gammaproteobacterium *Pseudoalteromonas luteoviolacea*. This bacterium has been found to synthesize structures, now known as tailocins, from a gene set gained from a T4-type bacteriophage [9,10]. These complex arrays of tailocins produced by inductive strains of *P. luteoviolacea* were originally dubbed metamorphosis-associated contractile systems, MACs [10]. However, neither arrays of tailocins nor their individual elements are found in some other inductive biofilm bacterial species, suggesting that other structures must be involved [11].

In a more recent paper by Shikuma et al. [11] focused on the early transcriptional events in larvae of *H*. elegans that have been cued to settle and metamorphosis in response to the tailocin arrays isolated in a crude preparation from *P. luteoviolacea*. The authors reported that the settlement process begins with loss of the prototrochal cilia that both propel and feed the larva [11]. This was in contrast to the report of Carpizo-Ituarte and Hadfield [1], who noted that settling larvae of *H*. elegans first tether themselves to a selected surface via a mucous thread, next secrete an encompassing primary tube attached to the surface, and then, enclosed in the primary tube, lose their motile cilia. Furthermore, larvae of *H. elegans* exist in a world of turbulent flow, even in bays and harbors. At our field site in Pearl Harbor, Hawaiʻi, a habitat for *H. elegans*, we measured flow rates of <6 cm s^−1^ and turbulence from wind chop and ship wakes [12]. This strongly suggests that ‘*anchor first, metamorphose second*,’ must be the rule for a settling larva to avoid being swept away after losing its motile cilia during metamorphosis. We have thus re-visited the question and compared the order of metamorphic events when competent larvae of *H. elegans* contact a naturally occurring biofilmed surface, a single-species biofilm of the inductive bacterium *P. luteoviolacea*, or are induced to settle by the semi-purified tailocin arrays from *P. luteoviolacea* described and labeled “MACs” by Shikuma et al. [10]. We visually followed and video-recorded the settlement and developmental events for 30 – 60 minutes and recorded the timing of each.

Because the observations made during the procedures described above suggested that the action of tailocin arrays was primarily destruction of cilia and ciliated cells, we performed two additional experiments. First, we exposed pre-competent larvae of *H. elegans* -- that is, larvae not yet developmentally capable of undergoing metamorphosis -- to the tailocin-array preparation to determine if these larvae might suffer destruction of their prototrochal cells or cilia. Secondly, we exposed competent veliger larvae of the coral-eating nudibranch gastropod *Phestilla sibogae*, who metamorphose only in response to a soluble cue from their prey coral, to the tailocin-array preparation to determine if the prominent ciliated cells of the velum, responsible for swimming and feeding, would be affected.

## Materials and Methods

### Culture of H. elegans

Adult worms were collected at our field site in Pearl Harbor, Hawaiʻi and maintained in running seawater at the Kewalo Marine Laboratory, Honolulu, HI (KML). We spawned adult worms and reared larvae of *H. elegans* according to the protocols described by Nedved and Hadfield [2]. Metamorphically competent larvae, 5 – 6 days postfertilization, were induced to settle by three different preparations: (1) a natural biofilm; (2) a monospecific biofilm of *P. luteoviolacea* (H1 strain); or (3) exposure to a suspension of the inductive tailocin arrays (MACs) of *P. luteoviolacea*.

### Experimental preparations

Natural biofilms were accumulated on glass microslides by submersion for three weeks or longer in aquaria at the KML supplied with continuously flowing, unfiltered seawater in either an open position exposed to ambient light (WT-BF) or in a black container without light (BB-BF). The biofilms formed in the latter lack the heavy coating of diatoms that develop on slides exposed in natural light. These slides were scored with a diamond tipped pen, broken, and a piece of the biofilmed slide (approx. 22 mm X 22 mm) was used as an inductive cue. Monospecific biofilms of *P. luteoviolacea* H1 were allowed to form on pieces of glass microslides by inoculating them with 10^7^ cells ml^−1^ from an overnight broth culture of *P. luteoviolacea* for 1 h [6]. The crude preparation of tailocin arrays and other cell products from *P. luteoviolacea* was prepared as per Shikuma et al. [10]. A 100-fold dilution in 0.22 μm double-filtered, autoclaved seawater (DFASW) of this preparation was used as the cue. *Treatment of competent larvae of H. elegans*. Mono- and multi-species biofilmed slide pieces were put onto a complete microscope slide for handling. An aliquot of 0.22 μm-filtered seawater (FSW) containing competent larvae of *H. elegans* was added to the top of the slide fragment. To administer the tailocin-array preparation, larvae were added to a 35 mm petri dish containing the diluted preparation, and an aliquot of this mixture was added to a clean microscope slide for observation.

### Treatment of pre-competent larvae of H. elegans and larvae of a nudibranch

We exposed 2-day old, pre-competent trochophore larvae of *H. elegans* and veliger larvae of the nudibranch gastropod *Phestilla sibogae* to the tailocin-array preparation and recorded the results as described below. Veliger larvae were obtained from a captive population of *P. sibogae* maintained at KML.

### Video-recording

After adding larvae to each of the above preparations, they were observed under a dissecting microscope to determine if they had begun the slow, circular swimming characteristic of pre-settlement behavior for the *Hydroides* larvae, or slow swimming while gliding across the surface, foot down, for the *Phestilla* larvae. When these behaviors were observed, a coverslip with supports at each corner was added to the slide, and it was transferred to the stage of a Zeiss microscope equipped for bright-field, phase-contrast and interference-contrast microscopy. The larvae were then observed, and their behavior video recorded with a camera (Canon Rebel T1i EOS 500D) mounted on the microscope. Additionally, the sequences, timing and durations of their behavioral events were manually recorded. Video recordings were made of larvae from ten different larval cultures over a five-month period. During recording, we particularly focused on the initial events involving larval attachment, cilia loss and primary-tube formation.

### Event sequence analysis

The settlement and metamorphosis events were analyzed using hierarchical clustering with the TraMineR package in R (version 2.2-0.1) [13, 14]. Event sequences were clustered using the Optimal Matching for the dissimilarity matrix [13–15]. This method focuses on the order of events rather than the time spent in each event. Samples are clustered based on similarity, with differences in the sequences receiving a gap penalty 1 and a single substitution cost of 2. Larval searching behavior, attachment, cilia loss, cell loss and primary tube formation were included in the analysis.

## Results

While timing of the onset of events varied from larva to larva, when larvae were exposed to a natural multispecies biofilm the order of events was always the same: (1) the larvae stop swimming by attaching to the substratum with a mucus strand (Fig. 2); (2) the larvae secrete a transparent primary tube which encases the entire larva, but remains open at the anterior end (Figs. 2 and 3a); (3) the larvae shed their prototroch cilia, and their collars deflect anteriorly (Figs. 2 and 3b); (4) the larvae shed their food groove cells and evert the toe-like neuropodia from the posterior edge of their third segment (Figs. 2 and 3c); (5) the larvae begin to remodel the episphere and develop branchial filaments. When larvae were exposed to a biofilm composed only of *P. luteoviolacea*, this sequence was also largely the same, with all larvae attaching and secreting primary tubes before loss of cilia (Fig. 2) (See Supporting Table ST1 and Supporting Video SV1). In both instances, shed cilia and cells are typically directed to the food groove toward and the mouth and swallowed (see Supporting Video SV1).

**Fig. 2.**
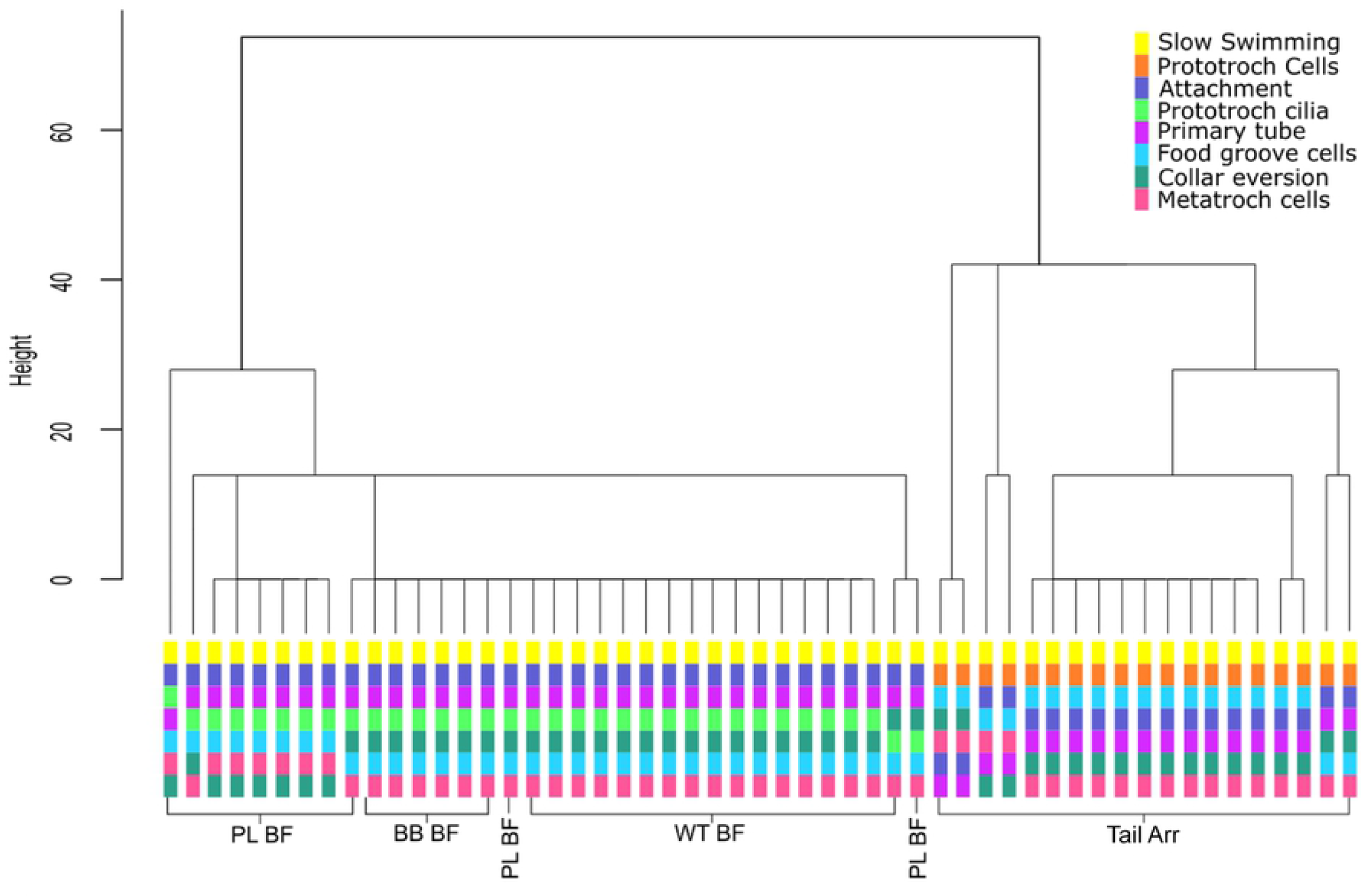
Dendrogram of Hierarchical Cluster Analysis using Optimal Matching generated with the TraMineR package [13]. The height of the vertical lines reflects the level of separation. Below the dendrogram, the sequence of events for each sample is included. Samples are labelled according to the settlement inducing treatment they received: Tail Arr, semi-purified preparation of tailocin arrays and cellular products from *P. luteoviolacea*; PL BF, biofilm of *P. luteoviolacea*; BB BF, natural biofilms accumulated in the absence of light; WT BF, natural biofilms accumulated in the presence of light.

**Fig. 3.**
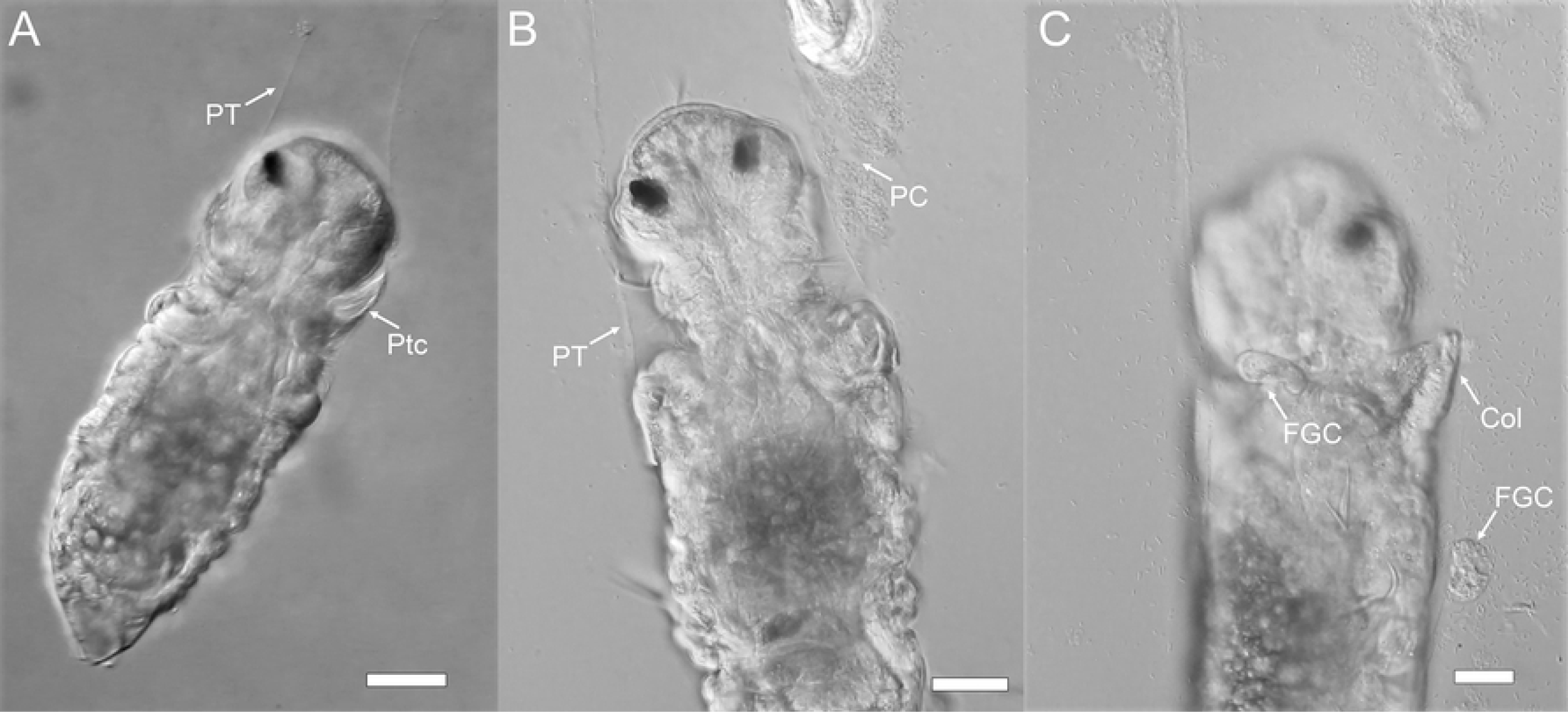
Sequence of metamorphic events in competent larvae of *H. elegans* exposed to a natural biofilm. A) prototroch cilia are still attached when primary tube begins to form.; B) Prototroch cilia shed while larvae is in primary tube; C) Loss of food-groove cells and collar eversion. Scale bar 20 μm. Ptc, Prototroch cells; PT, Primary Tube; PC, shed Prototroch cilia; Col, collar; FGC, food groove cells.

When larvae were exposed to the crude extract of tailocin arrays and other cell products from *P. luteoviolacea*, the sequence of events was highly variable but consisted mostly of the following sequence: (i) stopped swimming while secreting copious mucus which did not act as a tether; (ii) shed their prototroch cilia; (iii) shed their food-groove cells and prototroch cells (Figs. 2 and 4); and then, (iv) secreted a primary tube (Fig. 2)(Supporting Table ST1; Supporting Video SV2). In all instances of exposure to the bacterial products, cilia loss was observed before primary tube formation (Fig. 4). The entire contour of the larvae body became distorted during response to treatment with the tailocin-array preparation. Because larvae are not enclosed in a primary tube during loss of cilia and cells, these products, normally eaten during metamorphosis, simply float away,

**Fig. 4.**
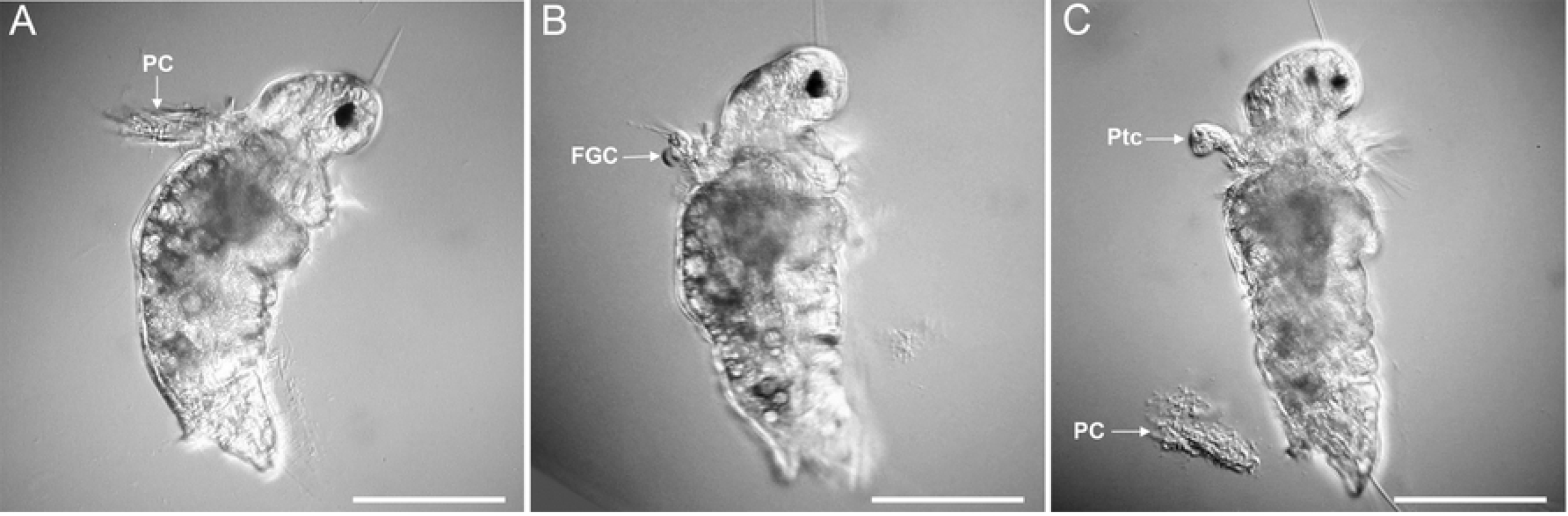
Larva of *H. elegans* induced to metamorphose by a tailocins-array preparation. (A) loss of prototroch cilia before attachment and primary-tube formation; (B) loss of food-groove cells before attachment and primary-tube formation; (C) loss of prototroch cells before attachment and primary-tube secretion. FGC, food-groove cells; PC, prototroch cilia; Ptc, prototroch cilia. Scale bars, 50 μm.

When pre-competent larvae of *H. elegans* were exposed to the crude preparation of tailocin arrays, they shed their prototrochal cells and cilia without undergoing any further stages of metamorphosis (Fig. 5) (Supporting Video SV3). And when the veliger larvae of the nudibranch *Phestilla sibogae* were similarly exposed, they also began to shed their large ciliated velar cells (Fig. 6) (Supporting Video SV4), which are homologues of the prototroch cells of other molluscs and polychaetes.

**Fig. 5.**
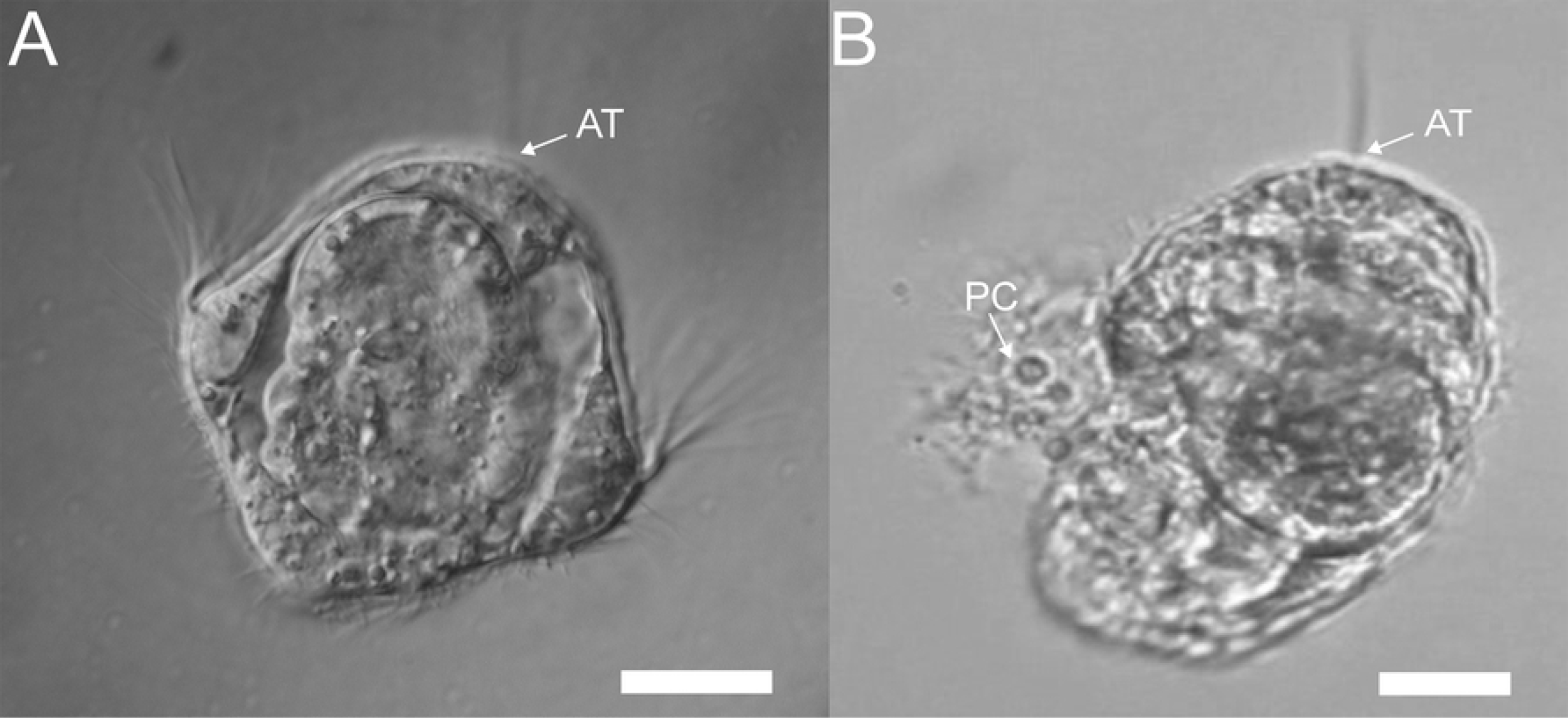
Pre-competent larvae of *H. elegans* lose cilia and prototroch cells when exposed to the semi-purified tailocin arrays from *P. luteoviolacea*. (A) Untreated pre-competent trochophore larva of *H. elegans*. (B) Treated pre-competent larvae of *H. elegans* showing loss of prototroch cells and cilia. AT, apical tuft of cilia; PC, prototroch cells detaching from the larva. Scale bar = 20 μm.

**Fig. 6.**
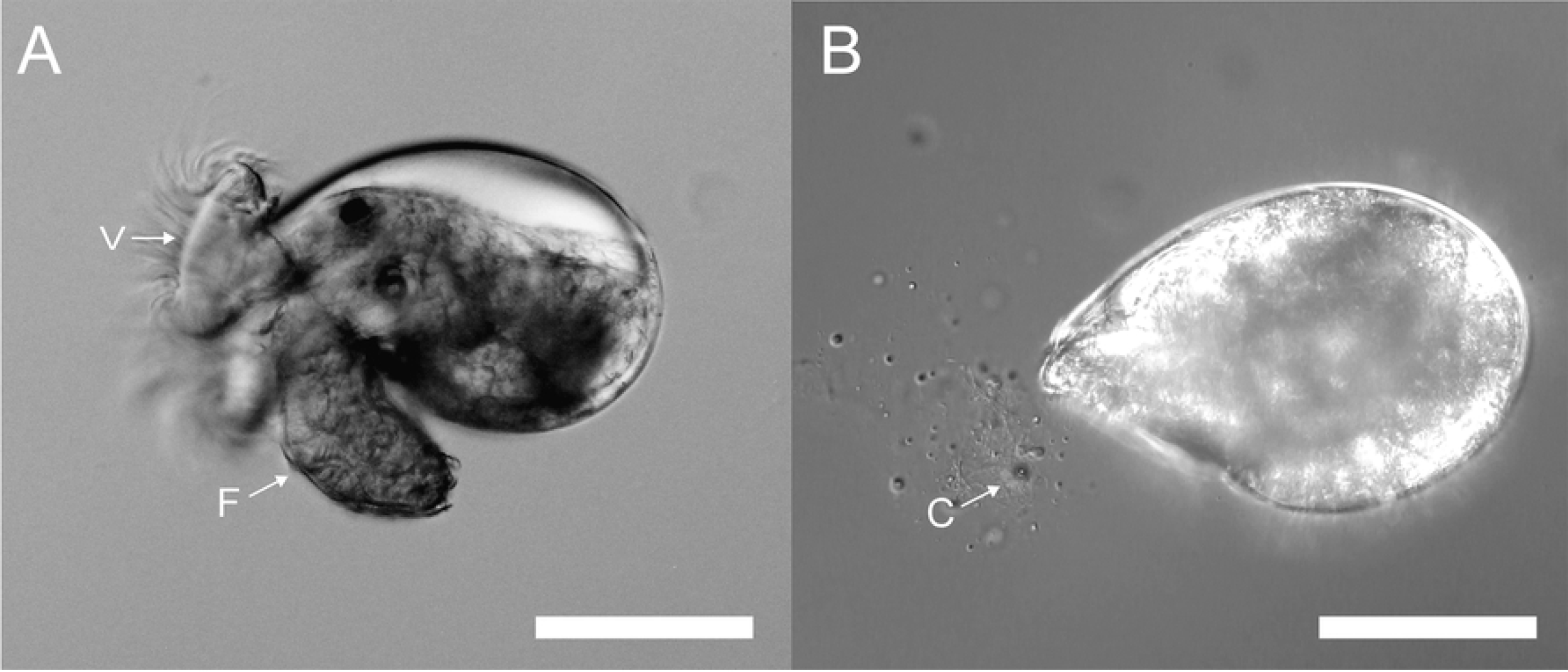
Larva of *P. sibogae* lose cilia when exposed to the semi-purified tailocin arrays from *P. luteoviolacea*. (A) Untreated larva from *P. sibogae* showing velar cilia; F: foot, V: velum. (B) Contracted larva exposed to tailocin arrays from *P. luteoviolacea* shed cilia and ciliated cells (C). Scale bar 100 μm.

## Discussion

An initial paper on the larval biology of *H. elegans* noted its dependence on marine biofilms to initiate settlement and metamorphosis [3]. Subsequently, many papers have appeared on this topic focusing on both the bacterial specificity in settlement and the developmental biology of this widely distributed marine worm. Among the latter, Carpizo-Ituarte and Hadfield [1] characterized the events of settlement and metamorphosis on wild-type biofilms, noting the same sequence found in the current investigation: (1) surface exploration; (2) attachment; (3) primary-tube formation; (4) prototroch loss; and (5) morphogenesis of the anterior region. Investigations by Walters et al. [16] and Koehl and Hadfield [12] focused on details of the hydrodynamic forces in habitats inhabited by *H. elegans*, which revealed that larvae of this species are always subject to turbulent, laminar flow when attempting to settle. The studies made it clear that forming a strong attachment to a surface before loss of motility is of paramout importance to the recruitment of *H*. elegans. Flow rates of 2 – 6 cm/sec, measured along surfaces Pearl Harbor, HI [12], would carry a larva a meter or more from a selected settlement site within a second, had it not firmly attached at that site. Because of this, we reason that secretion of the primary tube *before* loss of the larval swimming organ, the prototroch, is crucial to successful recruitment.

The difference in the order of metamorphic events induced by the crude preparation of tailocin arrays may also be simply a result of the experimental setup. The tailocin arrays of *P. luteoviolacea* are exceptionally delicate, and their activity can be destroyed by gentle filtration [10]. As a consequence, the current method of isolation uses low speed centrifugation to remove cells and produces a concentrated but complex solution which also contains outer membrane vesicles and other cellular materials [10, Fig. 2D]. It is also highly likely that the preparation contains secondary metabolites such as violacein and other lightweight cellular debris. This experimental difficulty necessitates that the larvae are exposed to the tailocin arrays in the form of a bath to demonstrate the tailocin bioactivity. Since it has been clearly demonstrated with naturally occurring biofilms that larvae of *H. elegans* must physically touch the biofilms to detect and respond to the metamorphic cue, the use of a bath exposure, although a helpful screening tool, must be recognized as not ecologically relevant. This situation is far from unusual in the chemical ecology field, but it does serve as a timely reminder that experimental results do not always reflect the ecological context of the setting where the phenomenon under study would naturally occur.

In the competent larva of *H*. elegans, there are a number of transcripts that are regulated during stimulation of metamorphosis by the tailocin arrays from *P. luteoviolacea* [11]. The p38/JNK MAPK signaling pathways that are associated with metamorphosis in the larva appear to be implicated in the morphogenic events associated with metamorphosis and not cilia loss induced by tailocins [11]. However, in addition to their potential roles in cell adhesion and innate immunity, a subset of these transcripts is also associated with other cellular processes including defense against pathogens [17,18,19] and inflammatory responses [20]. The up and downregulation of these transcripts may also be a response to the cellular insult to the ciliated cells of the trochal bands by the tailocin preparations. The revelation that the order of morphogenetic events during settlement and metamorphosis of larval *H. elegans* is badly disturbed when induced by the crude tailocin extract confounds our ability to ascertain the true involvement of the identified transcripts in naturally induced metamorphosis of larvae of *H. elegans*.

The additional observations reported here, that the crude preparation of tailocin arrays is destructive to larval ciliated cells in pre-competent larvae of *H. elegans* and unrelated larvae of a nudibranch strongly, suggest that the complexes have a natural function other than inducing settlement of specific polychaete larvae. Perhaps their evolutionary value lies in destroying ciliated predators when cells of *P. luteoviolacea* are resident in biofilms. Freckelton et al. reported settlement induction by other bacterial species that lack the genes for tailocin arrays [20] and that lipopolysaccharide isolated from inductive *Cellulophaga lytica*, which does not make tailocins, induces larvae of *H. elegans* to settle [21]. There is still much to be learned about the induction of settlement in *H. elegans* by surface films made by any single bacterial species, and especially by *Pseudoalteromonas luteoviolacea*.

## Acknowledgments

The authors gratefully acknowledge the excellent laboratory assistance of Ms. Amy Knowles.

## Supporting Information

**ST1 Table 1. Mean time and range, in minutes, to the start of the settlement and metamorphosis events in *Hydroides elegans*.**

**SV1 Video 1. Sequence of metamorphic morphogenesis initiated by biofilms.** A metamorphically competent larva of *Hydroides elegans* has been placed on a biofilm composed of the bacterium *Pseudoalteromonas luteoviolacea* and videotaped as the events of settlement/attachment and metamorphosis proceed. The observed events are the same when a larva induced to settle by a complex, wild-type biofilm, but less easy to observe due to a dense background of diatoms and cyanobacterial filaments. The primary tube is very transparent and can best be seen when it accumulates bacterial cells from the underlying biofilm. The larva moves constantly back and forth and rotates while secreting the primary tube. Typically, cilia from the prototroch and the cells of the food groove are swept into the food groove, then to the mouth and swallowed. FCG, food-groove cells; Muc, mucus; NP, neuropodia; PT, primary tube.

**SV2 Video 2. Sequence of metamorphic morphogenesis initiated by tailocin arrays.** A metamorphically competent larva of *Hydroides elegans* has been exposed to a suspension that includes the crudely separated tailocin arrays produced by the bacterium *Pseudoalteromonas luteoviolacea*, known to induce metamorphosis (see Shikuma et al. 2014). Without first attaching, the larva sheds prototroch cilia, food groove cells and prototroch cells, and subsequently produces a primary tube. Because the larva is not enclosed in a primary tube, shed cilia and cells float away; they are typically eaten by a larva during normal metamorphosis. FGC, food-groove cells; PC, prototroch cilia; PT, primary tube; Ptc, prototroch cells.

**SV3 Video 3. Tailocin arrays trigger loss of cilia and cells in precompetent larvae.** A trochophore larva of *Hydroides elegans*, developmentally incapable of undergoing metamorphosis, has been exposed to a suspension that includes the crudely separated tailocin arrays produced by the bacterium *Pseudoalteromonas luteoviolacea*, known to induce metamorphosis in competent larvae. The larva sheds both ciliated cells of its prototroch and from its food groove. None of the larvae treated with the tailocins-array preparation underwent metamorphosis. FGC, food-groove cell.

**SV4 Video 4. Tailocin arrays trigger loss of cilia and cells in veliger larvae.** A metamorphically competent veliger larva of the nudibranch gastropod *Phestilla sibogae* has been exposed to a suspension that includes the crudely separated tailocin arrays produced by the bacterium *Pseudoalteromonas luteoviolacea*, known to induce metamorphosis in larvae of *Hydroides elegans*. The larva retracts and secretes mucus from its foot, then begins to lose ciliated cells from either its foot or velum. Soon after, the larva begins to shed the large multi-ciliated cells of its velum. Retained for an additional 24 hours, none of the treated larvae metamorphosed. VC, velar cell.

## References

1. Carpizo-Ituarte E, Hadfield MG. Stimulation of metamorphosis in the polychaete *Hydroides elegans* Haswell (Serpulidae). Biol Bull. 1998;194: 14–24.

2. Nedved BT, Hadfield MG. *Hydroides* elegans (Annelida: Polychaeta): A Model for Biofouling Research. Marine and Industrial Biofouling. 2008. pp. 203–217. doi:10.1007/7142

3. Hadfield MG, Unabia CC, Smith CM, Michael TM. Settlement preferences of the ubiquitous fouler *Hydroides elegans*. Recent developments in biofouling control. 1994. pp. 65–74.

4. Unabia CRC, Hadfield MG. Role of bacteria in larval settlement and metamorphosis of the polychaete *Hydroides elegans*. Mar Biol. 1999;133: 55–64. doi:10.1007/s002270050442

5. Lau SCK, Qian PY. Larval settlement in the serpulid polychaete *Hydroides elegans* in respouse to bacterial films: An investigation of the nature of putative larval settlement cue. Mar Biol. 2001;138: 321–328. doi:10.1007/s002270000453

6. Huang S, Hadfield MG. Composition and density of bacterial biofilms determine larval settlement of the polychaete *Hydroides elegans*. Mar Ecol Prog Ser. 2003;260: 161–172. doi:10.3354/meps260161

7. Vijayan N, Hadfield MG. The microbial diversity of a marine biofilm that induces larvae of *Hydroides elegans* (Polychaeta) to settle and metamorphose. Environ Microbiol. 2020.

8. Hadfield MG, Nedved BT, Wilbur S, Koehl MAR. Biofilm cue for larval settlement in *Hydroides elegans* (Polychaeta): is contact necessary? Mar Biol. 2014;161: 2577–2587. doi:10.1007/s00227-014-2529-0

9. Huang Y, Callahan S, Hadfield MG. Recruitment in the sea: Bacterial genes required for inducing larval settlement in a polychaete worm. Sci Rep. 2012;2: 228. doi:10.1038/srep00228

10. Shikuma NJ, Pilhofer M, Weiss GL, Hadfield MG, Jensen GJ, Newman DK. Marine tubeworm metamorphosis induced by arrays of bacterial phage tail-like structures. Science (80-). 2014;343: 529–533. doi:10.1126/science.1246794

11. Shikuma NJ, Antoshechkin I, Medeiros JM, Pilhofer M, Newman DK. Stepwise metamorphosis of the tubeworm *Hydroides elegans* is mediated by a bacterial inducer and MAPK signaling. Proc Natl Acad Sci. 2016;113: 10097–10102. doi:10.1073/pnas.1603142113

12. Koehl MAR, Hadfield MG. Hydrodynamics of Larval Settlement from a Larva’s Point of View. Integr Comp Biol. 2010;50: 539–551.

13. Gabadinho A, Ritschard G, Müller NS, Studer M. Analyzing and Visualizing State Sequences in R with TraMineR. J Stat Softw. 2011;40: 1–37.

14. Gabadinho A, Ritschard G, Studer M, Müller NS. Mining Sequence Data in R with the TraMineR package: A user’s guide. Department of Econometrics and Laboratory of Demography, University of Geneva. 2009.

15. Studer M, Ritschard G. “What matters in differences between life trajectories: A comparative review of sequence dissimilarity measures”,. J R Stat Soc Ser A. 2016;179: 481–511.

16. Walters L, Hadfield M, del Carmen K. The importance of larval choice and hydrodynamics in creating aggregations of *Hydroides elegans* (Polychaeta: Serpulidae). Invert Biol. 1997;116: 102–14.

17. Donpudsa S, Tassanakajon A, Rimphanitchayakit V. Domain inhibitory and bacteriostatic activities of the five-domain Kazal-type serine proteinase inhibitor from black tiger shrimp *Penaeus monodon*. Dev Comp Immunol. 2009;33: 481–8.

18. Ward S, O’Sullivan J, O’Donnell J. von Willebrand factor sialylation—A critical regulator of biological function. J Thromb Haemost. 2019;17: 1018–1029. doi:https://doi.org/10.1111/jth.14471

19. Lenting PJ, Casari C, Christophe OD, Denis CV. von Willebrand factor: the old, the new and the unknown. J Thromb Haemost. 2012; 2428–2437. doi:https://doi.org/10.1111/jth.12008

20. Freckelton ML, Nedved BT, Hadfield MG. Induction of Invertebrate Larval Settlement; Different Bacteria, Different Mechanisms? Sci Rep. 2017;7. doi:10.1038/srep42557

21. Freckelton ML, Nedved BT, Cai Y-S, Cao S, Turano H, Alegado RA, Hadfield MG. 2020. Bacterial lipopolysaccharide induces settlement and metamorphosis in a marine larva. bioRχiv. doi.org/10.1101/851519

